# An invariant schema emerges within a neural network during hierarchical learning of visual boundaries

**DOI:** 10.1101/2025.01.30.635821

**Authors:** James R. Elder, Jie Zheng, Lydia B. Shimelis, Ueli Rutishauser, Milo M. Lin

## Abstract

Neural circuits must balance plasticity and stability to enable continual learning without catastrophic forgetting, a pervasive feature of artificial neural networks trained using end-to-end learning (e.g. backpropagation). Here, we apply an alternative, hierarchical learning algorithm to the cognitive task of boundary detection in video clips. In contrast to backpropagation, hierarchical training converges to a network executing a fixed schema and generates firing statistics consistent with single-neuron recordings from human subjects performing the same task. The hierarchically trained network’s schema circuit remains invariant following training on sparse data, with additional data serving to refine the upstream representation.

---

How neural networks solve cognitive tasks in a way that prioritizes model robustness while allowing plasticity is an open problem (*1–3*). Artificial neural networks (ANNs) are susceptible to catastrophic forgetting, wherein learning from additional data overwrites system parameters (*4, 5*), whereas the brain is capable of continuous learning and model refinement while retaining previously learned core cognitive structures (*6, 7*)–referred to as “schemas” in cognitive psychology (*8*).

Neural circuits can balance robustness and plasticity by creating subpopulations of stable and plastic neural connectivities. In vision, previously plastic sensory thalamic inputs in the visual cortex are stabilized in adulthood, while higher-order visual representations remain plastic (*9–12*). Similarly, auditory fear conditioning suggests that synapse-specific plasticity encodes memory associations, whereas engram cells form a stable network, allowing specific memory erasure without schema disruption (*13*). However, the self-organizing principles by which neural network connectivities are trained to produce schemas that are sensitive to–but whose cognitive structure is robust to–new inputs are unknown (*14*).

Longstanding limitations in spatial and temporal resolution preclude a purely experimental solution to this problem. Computational modeling can potentially identify general mechanistic principles (*15–17*) and make testable predictions about connectivity, perturbation response, and firing patterns (*18–23*). Until recently, ANNs capable of learning general-purpose input-output tasks at scale have been trained using an **end-to-end** learning framework where the error of the network output is minimized by parameter adjustments propagated back through the network (*24–26*). Initialized with random parameters, the **Backpropagation** algorithm minimizes the task-specific error via gradient descent in parameter space. In this approach, neural connectivities and couplings are meaningful only insofar that they decrease output error.

Recently, we introduced a general-purpose **hierarchical** learning model that intrinsically produces neurobiological circuit properties missing from backpropagation trained networks (*27–29*). Neurons in these Essence Neural Networks (**ENNs**) are trained to perform stand-alone pair-wise distinguishing tasks and then hierarchically combined to perform the circuit task. This layer-wise learning approach increases network interpretability, enabling ENNs to be automatically *distilled* into equivalent human-readable code while demonstrating superior performance to backpropagation across an array of generalization benchmarks (*30*).

Here, we ask if networks trained using these two alternate learning paradigms on a cognitive vision task are differentially susceptible to catastrophic forgetting, here defined as the absence of subcircuits with invariant connectivities throughout training. Additionally, what are the implications of catastrophic forgetting on model performance, generalizability, robustness to noise, and statistical signatures of neuronal firing? Finally, do biological circuits display characteristic signatures of either training approach? We address these questions for feed-forward neural network architectures in the context of an image-based boundary-detection task.

Human experience is continuous, comprised of diverse internal and external neuronal and sensory inputs. However, memory encoding requires discrete segmentation of beginnings and endings (*31, 32*). Therefore, proper recognition of **boundaries** separating distinct visual inputs is required for effective memory encoding. In recent work (*33*), we identified two distinct neuronal cell types, **Boundary** and **Event** cells, in single-cell neuron recordings in the medial temporal lobe (**MTL, Fig. 1c**) implicated in boundary detection (**Fig. 1a**). Boundary cells fire shortly after the onset of a boundary (seen in Soft and Hard Boundary clips), whereas Event cells fire only for Hard boundaries approximately 100 ms after Boundary cells (**Fig. 2a-c**). Here, we trained both end-to-end and hierarchical neural networks on this visual boundary-detection task.

**Figure 1:**
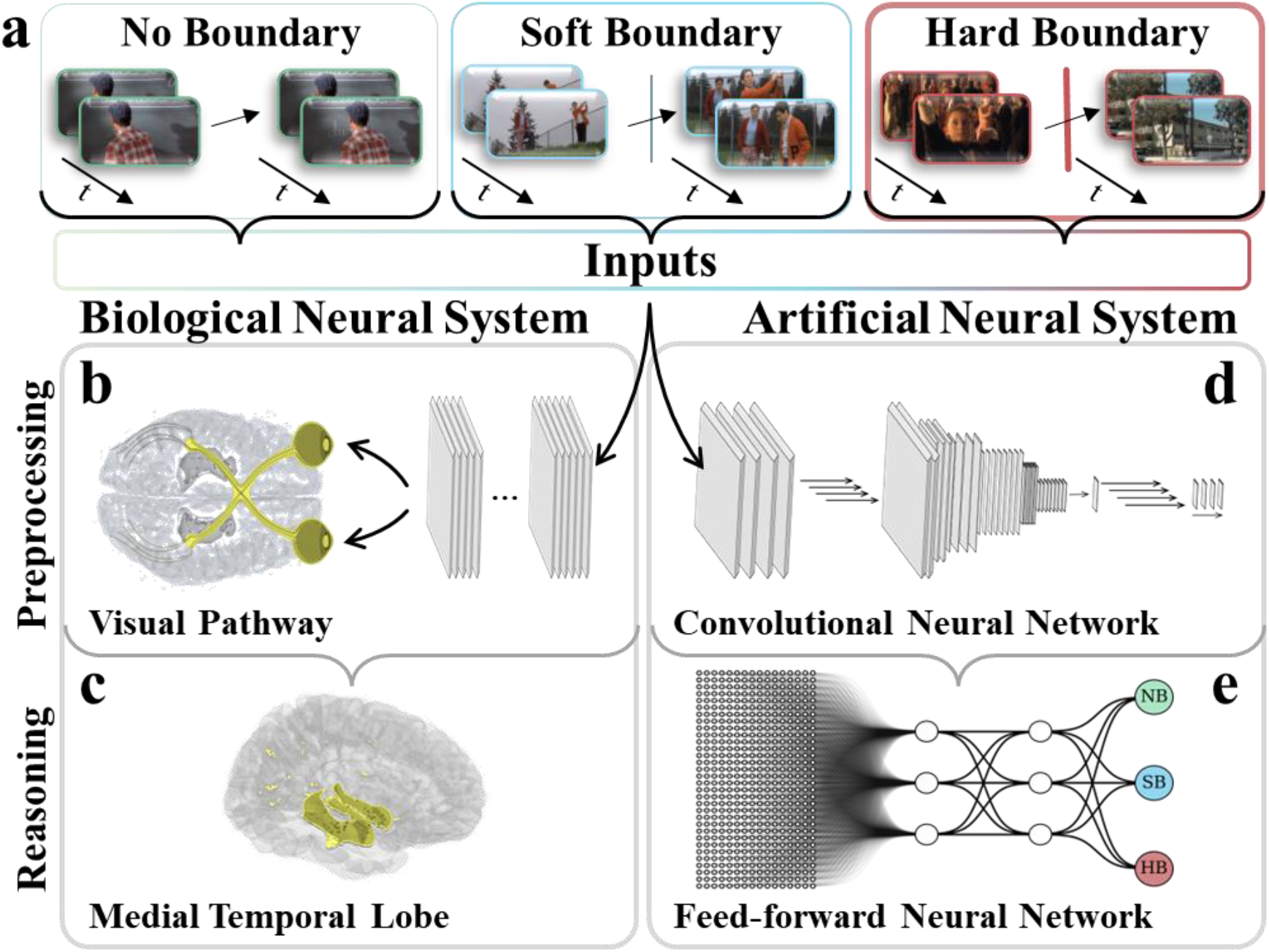
Boundary detection in video clips. **a**) Video clips containing either No Boundary, a Soft Boundary, or a Hard Boundary make up the inputs to both the biological neural system (*left*) and the artificial neural system (*right*). **b**) The visual pathway relays information from the retina to the visual cortex of the occipital lobe. **c**) Deep brain structures, such as the medial temporal lobe, interpret the processed visual stimuli. In the artificial neural systems (*right*), representative frames from the video clips are passed through a pretrained convolutional neural network (**d**) to generate the features fed into the cognitive top model (**e**) for training and evaluation.

**Figure 2:**
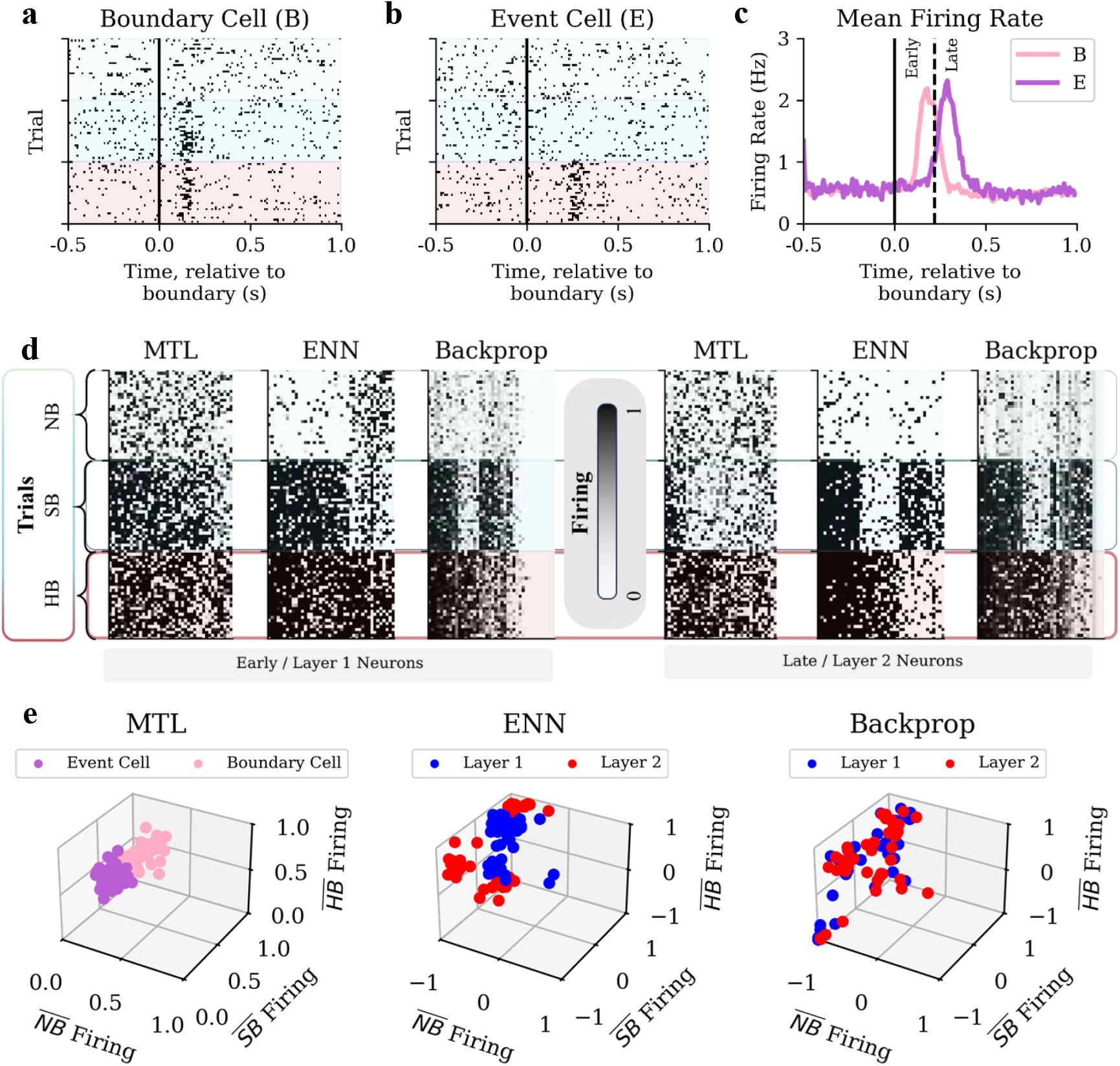
Neuron firing in boundary-detecting systems. **a**) Example MTL recordings of a Boundary cell and (**b**) an Event cell. Each neuron’s firing across an aligned 1.5 second time interval of 10 ms bins is shown for 90 different trials, 30 for each class (NB-green, SB-blue, and HB-red). **c**) Average boundaryaligned firing rate of Boundary (on all SB and HB trials, 42 cells, 105 trials) and Event (on all HB trials, 36 cells, 30 trials) cells. Cells are classified as “early” or “late” firers based on where they occur relative to the cutoff 220 ms after the boundary, roughly where the averaged peaks cross. **d**) Firing of early/hidden layer 1 neurons (*left*) and late/hidden layer 2 neurons (*right*). MTL cells (41 early, 37 late) are averaged over their respective peak firing windows (average peak across respective cell types ±2 st.d.), then activated through a sigmoid with steepness proportional to the background firing. Neurons from converged ENN and Backprop networks trained on 20 varying 50-50% splits of the dataset shown. Average firing of neurons in *d* by class on 30 clips from each class, colored by neuron type (Event or Boundary cell for MTL) or location (Layer 1 or 2 for ENN/Backprop).

We show that the hierarchical ENN learning algorithm reproducibly converges to a stable schema that contains the Boundary and Event cells observed in neural recordings, even with sparse training. Further training data refines the input representation into the invariant schema, increasing accuracy. In contrast, backpropagation-based networks are continually reconfigured during learning, with no fixed schema. Consequently, ENNs display increased robustness to parameter noise and are distillable into a discrete circuit that offers a causal explanation for the experimental temporal delay between Boundary and Event cells. Together, these results suggest hierarchical learning can generically create experimentally consistent neural networks with cognitive reasoning structures that avoid catastrophic forgetting.

To model the pre-processing in the optic pathway (**Fig. 1b**), we transformed the pixel inputs into the visual embedding of a convolutional neural network (CNN), a successful model of object recognition in the cortex (*34, 35*). For example, CNNs unconstrained by neural data contain layers highly predictive of neural responses in both V4 and inferior temporal cortex when trained on naturalistic images (*36*). Representative frames from video clips shown to patients during neural recording (*33*) were fed through a CNN pre-trained on the ImageNet dataset (VGG-16, **Fig. 1d**) (*37, 38*). The concatenated embeddings were used to train the feed-forward boundary classification neural network (**Fig. 1e**). The network was trained using either Backpropagation (end-to-end) or the ENN learning framework (hierarchical) with an 85-15% train-test split on 978 video clips. Across multiple splits, converged networks with 100% training accuracy achieved an average test accuracy of 62.3% (±3.8%) and 56.1% (±4.7%) for ENNs and Backprop, respectively (**Sup. Fig. 1**), a roughly two-fold improvement over chance, with error primarily coming from distinguishing between soft versus hard boundary cases (**Sup. Fig. 2**). For the purpose of comparing hierarchical versus end-to-end learning, we will show that this level of performance is sufficient due to ENN schema fixation under sparse training (**Fig. 3c**).

**Figure 3:**
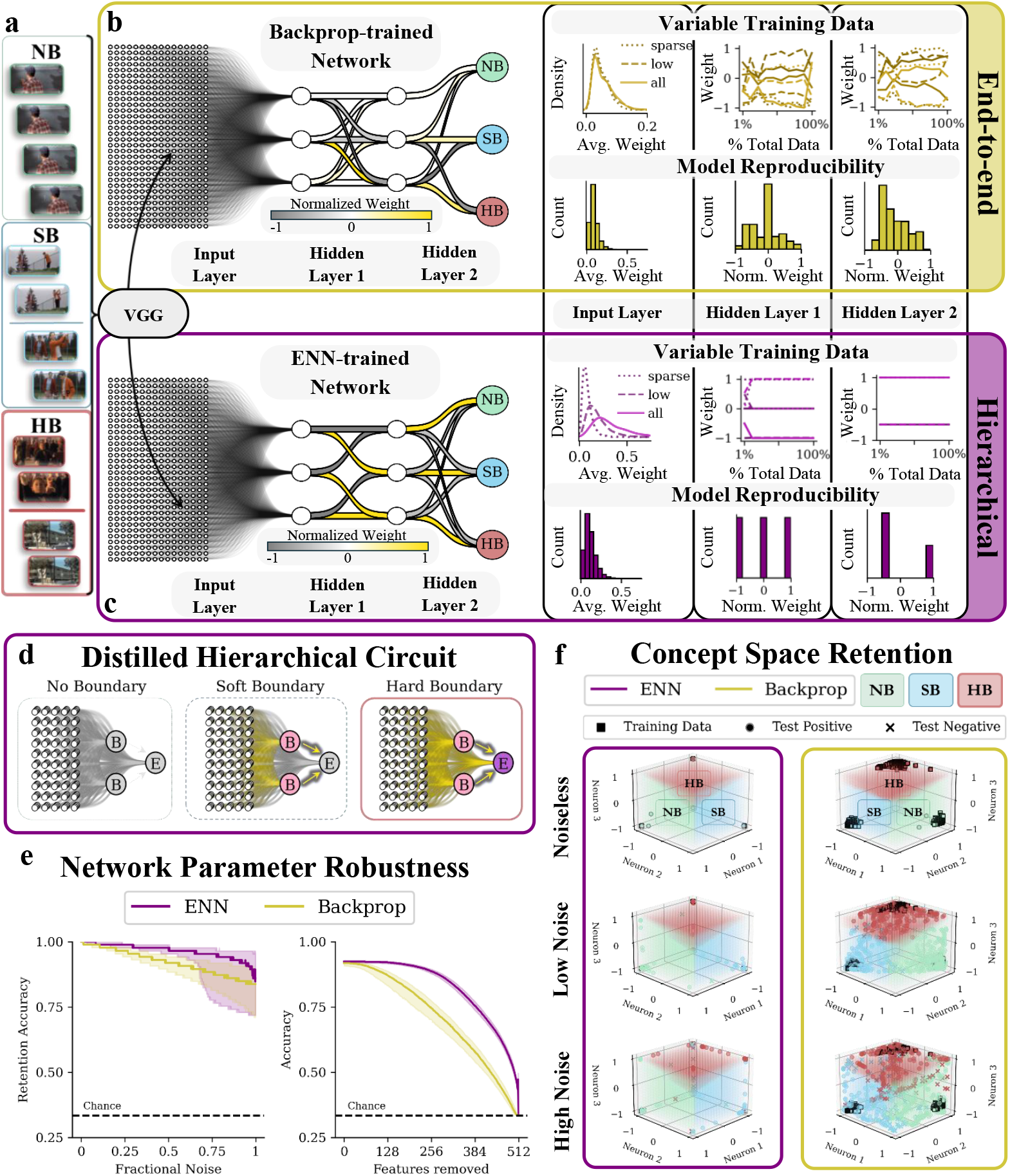
Artificial boundary-detecting neural systems. **a**) Representative frames (4 frames ×(1080 ×1920) pixels × 3 channels) are fed independently through VGG-16 and concatenated to serve as the input for the top models (4 frames 5×12 features). Example structure and learning trajectory (*Variable Training Data*) of a network trained using either end-to-end (**b**) or hierarchical (**c**) learning. Internal network weights colored by normalized weight. *Model Reproducibili ty*: Network weight distribution of converged networks from 10000 85-15% train-test splits. **d**) Distilled circuit firing by class. Top 50 features from the ENN are inputs to the circuit, while the boundary detection task is performed by Boundary and Event cells, sequentially. **e**) *left*, noise, sampled from a Gaussian distribution with standard deviation scaled to the learned weight, is injected into the weights between the 1st and 2nd hidden layers. *right*, features, ranked by importance, are replaced with their average input across all frames in the dataset. **f**) Hidden layer 2 concept space retention with noise injection: (*top*) unnoised network, (*middle*) lightly noised networks, (*bottom*) heavily noised networks. Each axis corresponds to the firing of a neuron on the 2nd hidden layer.

While boundary- and event-like firings were observed sporadically in networks trained with end-to-end learning (Backprop), their presence was strongly dependent on initialization conditions and displayed no statistical association between network layer and firing classes (**Fig. 2d-e**). In contrast, we observed that even sparsely trained ENNs reproducibly contained neurons whose firing patterns and ordering (**Fig. 2d**) were consistent with those from single-cell recordings (**Fig. 2b-c**). ENN and biological neurons, when characterized by class-specific firing patterns, were clustered and separable by layer, unlike backprop-trained neurons (**Fig. 2e**). Boundary-like neurons in ENNs (firing primarily on SB and HB clips) consistently appeared most prominently in the first hidden layer, while event-like neurons (firing on HB clips only) were enriched in the second hidden layer (**Fig. 2d-e**). This sequential ordering mirrors the temporal ordering of Boundary and Event cells observed experimentally (**Fig. 2c**). We distilled the ENN’s learned schema into a binary circuit that operates on only 50 features per frame (**Fig. 3d**, example features in **Sup. Fig. 4**), with only a marginal loss in performance (comparative performance of alternate models on the same features in **Sup. Fig. 5**). In the distilled circuit, the event-like cell is directly gated by the boundary-like cells (**Fig. 3d**), suggesting that the experimentally observed delay is due to the Boundary cells being upstream of Event cells.

Notably, hierarchical learning converges to its stable schema with limited training data (hidden layers, **Fig. 3c**, *Variable Training Data*). Additional data solely refines input layer weights, optimizing feature representation into the invariant schema and leading to stratification of feature importance (input layer, **Fig. 3c**, *Variable Training Data*). In contrast, networks with end-to-end learning (Backprop) displayed catastrophic forgetting, characterized by significant parameter heterogeneity (**Fig. 3b**, *Model Reproducibility*), reflecting both their random instantiation and unpredictable learning trajectories with substantial drift in all layers during training. Specifically, weights in the “cognitive” layers downstream of the input layer fluctuate without dampening when trained on increasing amounts of data (hidden layers, **Fig. 3b**, *Variable Training Data*). Interestingly, in contrast to the corresponding layer of the ENN, the input layer weight distribution of the Backprop-trained network does not change with increasing training data, indicating a lack of representational sparsification (input layer, **Fig. 3b**, *Variable Training Data*).

We next investigated the implications of these differences on robustness to fluctuations in synaptic strength. We directly perturbed the trained networks’ internal weights connecting the first two hidden layers. Upon adding random noise to the weights, the ENN retained its superior performance even at noise levels of the magnitude of the learned weights (**Fig. 3e**, *left*). Furthermore, the failure modes of these networks were qualitatively different: misclassifications induced by noise in the ENNs primarily occurred in a specific class, whereas Backprop networks displayed varied failure modes, likely due to their distributed computation (**Fig. 3f, Sup. Fig. 3**).

Previously, we showed that ENNs intrinsically possess neurobiological properties, such as modular connectivity and localized firing, that are missing in networks trained using backpropagation without additional priors/constraints (*27*). Yet, how the firing statistics of either model compares to that of biological neurons implicated in the same cognitive task was not addressed. In this study, for the visual boundary detection task, we demonstrate that only ENN networks consistently produce boundary and event cells whose firing statistics are consistent with the task separability and temporal ordering observed in neural recording data. We show that even during the early learning stage, hierarchical learning avoids catastrophic forgetting by refining the input representation while leaving the core schema invariant. Schema invariance despite overall network plasticity during early learning is a consequence of hierarchical learning and is distinct from schemas associated with accelerated transfer learning between related tasks reported in Recurrent Neural Networks trained on abstracted sensorimotor and serial learning tasks (*39, 40*). In contrast, end-to-end learning displays catastrophic forgetting, evidenced by marked sensitivity of the entire network to initialization conditions and data exposure. In the ENN, schema fixation during early learning arises because each sub-concept neuron computes an effectively digital “and” operation on its upstream differentia neurons which are continuously refined. This allows the schema to be mapped to an equivalent to digital circuit, and also makes the ENN network more robust to fluctuations in synaptic strength. Considering the task-agnostic nature of these properties, our results suggest that hierarchical learning is a general mechanism that allows invariant cognition structures to arise from distributed neural computation.

## Methods

### Dataset

The dataset consisted of 888 class balanced clips (298 NB, 293 SB, and 297 HB) selected analogously to and supplemented to those from the original experimental dataset (30 clips for NB and HB, 60 for SB) from Zheng et al. (*33*). Clips were labeled as either: No Boundary (a continuous shot with no cuts or transitions), Soft Boundary (a continuous scene with a non-disruptive transition, e.g., cut to another perspective of the same scene), or Hard Boundary (two clips from unrelated scenes artificially spliced together). A 85-15% train-test split was used for all analyses unless stated otherwise.

### Video embedding

While the authors expected that pre-trained video transformers might yield the richest representation, after assessing preliminary performance of representations from pre-trained models such as ViViT (*41*), we found that the highest accuracy for this task came from using a representation from a VGG-16 trained on ImageNet (*37*).

#### VGG-16

A modified VGG-16 model pretrained on ImageNet from *keras v3*.*5*.*0* was used to generate all embeddings for downstream classification. The input to the model is a 1080x1920x3 image, preprocessed by keras’s *preprocess input* function, which converts the images from RGB to BGR, zero-centers each color channel with respect to the ImageNet dataset, without scaling. The 3 fully-connected layers at the top of the network used for ImageNet classification were replaced (*include top=False*) by a GlobalAveragePooling2D layer, reducing the dimensionality of the embedding to the final 512 features used for the final representation for each frame used in the subsequent top model.

#### Frames

Four frames were sampled from the video clips: the first frame, the frame prior to the boundary (or at the midpoint for No Boundary clips), the frame subsequent to the boundary (or immediately after the midpoint for No Boundary clips), and the final frame in the clip. Frames were preprocessed using keras’s *vgg16*.*preprocces input* function described above. Each frame was passed independently through the modified VGG-16, yielding a final embedding of dimensionality **4x512** (4 frames, each with 512 features).

### Top Models

All top models take the same flattened **4x512** embedding from the modified VGG and are tasked with learning to predicting the class of each input (i.e., NB, SB, or HB). Models discussed here are explicitly identical in architecture (i.e., number of hidden layers = 2, number of interneurons on each layer = 3, activation function = *tanh*) and perform the same classical feed-forward neural network operations during inference (*y* = *tanh*(**w** · **x** + *b*)). The final classification is determine by *argmax(y*), where *y* is the output of the final layer.

#### Essence Neural Networks

Previously, we introduced a class of artificial neural networks called Essence Neural Networks (ENNs), which are trained using a non-gradient-based schema based on neuro-cognitive models. Our implementation follows that outlined in the original paper (*27*). Briefly, training samples are grouped by label, then unsupervised learning divides concepts into subconcepts. Subsequently, individual neurons on the first hidden layer (termed differentia neurons) are learned (i.e., weights and biases) by a linear support vector machine (SVM) that distinguishes between all pairs of subconcepts. These neurons are then used to train the second hidden layer neurons (termed subconcept neurons) through computation of a SVM on the outputs of the differentia neurons for a given subconcept and all outside concepts. Finally, the output of the subconcept neurons are used to compute an SVM between each concept and all training data belonging to other concepts. On all layers, a neuron’s unactivated output is given by *y* = **w** · **x** + *b*, where **w** is the learned weights, **x** is the inputs from the previous layer, and *b* is the learned bias. A *tanh* activation function is used for all models discussed. For all other hyperparameters (e.g., LearnSubconcept options), defaults were used. Here, for all models discussed, the final architecture of the network consists of 2 hidden layers (differentia and subconcept), each with 3 interneurons.

#### Backpropagation Networks

Backpropagation networks were training using *scikit-learn 1*.*5*.*2* MLPClassifier class of the neural network module. Briefly, the classifier trains iteratively at each step (*max iter: 10000*), updating the model parameters according to the partial derivatives of the loss function (*solver: ‘adam’*). During inference, unactivated neuron outputs are given by *y* = **w** · **x** + *b*, where **w** is the learned weights, **x** is the inputs from the previous layer, and *b* is the learned bias. A *tanh* activation function is used for all models discussed. Here, for all models discussed, the architecture of the network consists of 2 hidden layers (identical to ENNs), each with 3 interneurons. For all other hyperparameters (e.g., *learning rate init, regularization strength*), defaults were used.

### Trial Firing

Biological neurons firings were normalized with respect to their background firing (i.e., firing before the boundary). Subsequently, their were activated firings were calculated using a sigmoid of the form (*y* = *a* ∗ tanh(b(x − c)) + d), where *a* is a scaling factor of 0.5, *b* is the steepness, inversely proportional to the standard deviation of the background firing, *c* is the xOffset = 1.5, and *d* is the yOffset = 0.5. Biological neurons were classified as “early” or “late” based on whether their peak firing occurred before or after 220 ms (relative to the boundary), as shown in **Fig. 2c**.

Artificial neuron firings were taken from the activated firings of hidden neurons from networks trained with a 50-50% split during a forward pass on randomly selected clips from the entire data set (30 trials for each class).

### Circuit Distilling

The top 50 features with the largest decrease in performance upon perturbation from an ENN trained with access to the entire dataset were used as the input for the distilled circuit. The weights and biases from the fully trained ENN are used to connect the inputs to the discrete neurons. Downstream firing is binarized to all-or-none (*y >*= 0, where *y* = **w** · **x** + *b*), with the “pink” boundary-like neurons operating as an and gate for the “purple” event-like cell (i.e., the event-like cell’s firing is 0 unless both boundary-like cells fire).

### Network Robustness Analysis

#### Noise Injection Analysis

Random noise was injected into the internal weights of the top model (specifically, all the weights between the first and second hidden layers). The noise was sampled from a Gaussian distribution and scaled to the magnitude of the weights to which it is added. Retention accuracy was calculated for each noised network by evaluating it on clips correctly classified by the unnoised network.

#### Feature Sparsification Analysis

Ranked features were selected and replaced with their average value across the dataset one at at time across all frames. Subsequently, the models evaluated on the sparse dataset until all 512 features had been removed.

## Acknowledgments

We thank Brad Pfeiffer for providing helpful feedback on the manuscript. This work was supported by the National Institutes of Health grants R01GM125748 (M.M.L.), R00NS126233 (J.Z.), and T32 5T32GM131963 (J.R.E). This research was supported in part by the computational resources provided by the BioHPC supercomputing facility located in the Lyda Hill Department of Bioinformatics, UT Southwestern Medical Center.

## Author contributions

J.R.E and M.M.L. conceived the work, J.R.E. J.Z. and L.B.S. curated and formatted video clips. J.R.E. performed neural network modeling and analysis. J.R.E. and M.M.L. wrote the paper. J.Z., U.R., and M.M.L. supervised the work.

## Competing interests

Authors declare that they have no competing interests.

## Data and materials availability

Code and data used for calculations are publicly available on GitHub (https://github.com/jre411/bdENN).

## Supplementary Information

**Figure S1:**
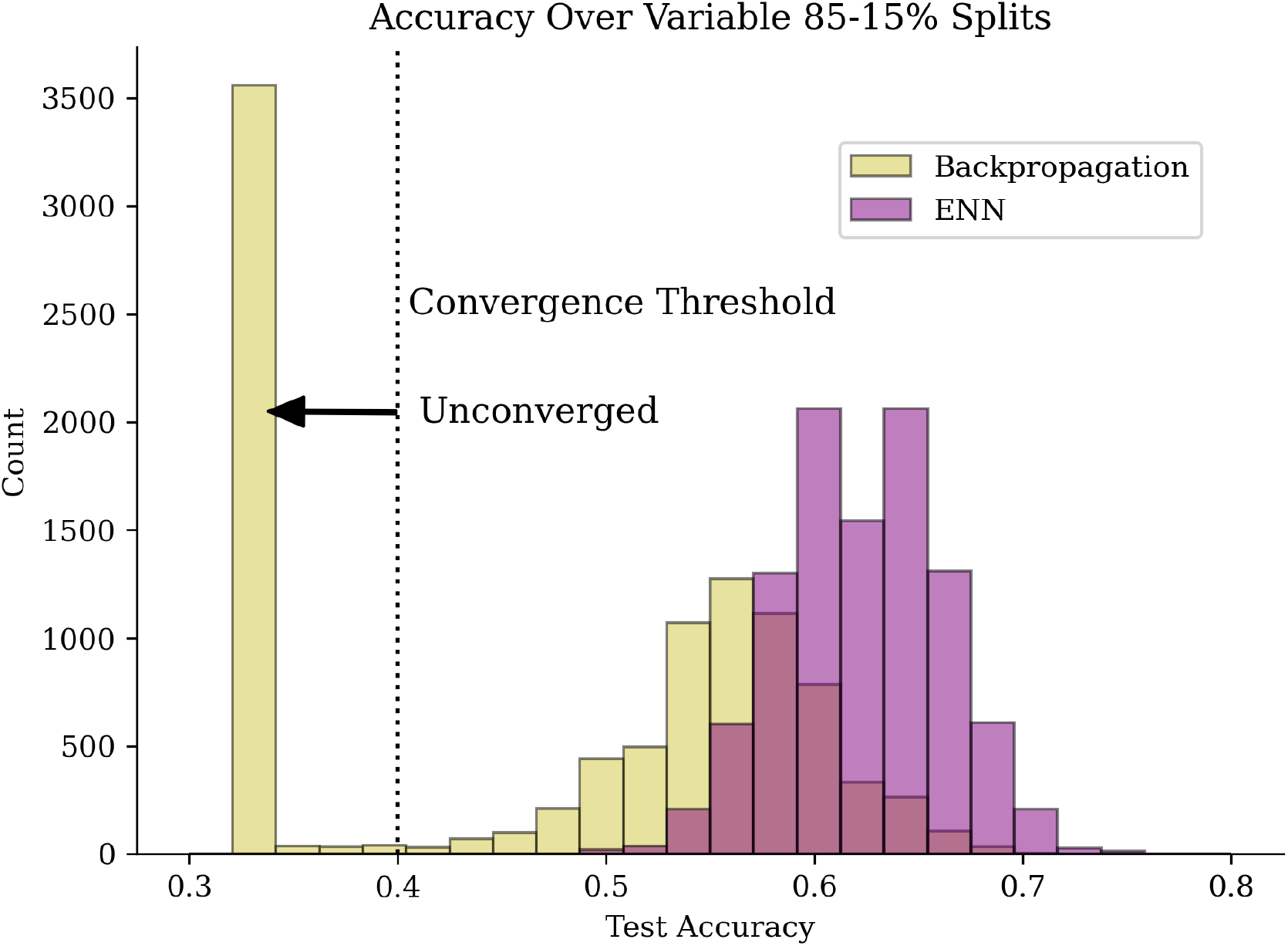
Boundary-detecting Performance. Histogram of network accuracies for Backpropagation and ENNs on variable 85-15% splits of the dataset.

**Figure S2:**
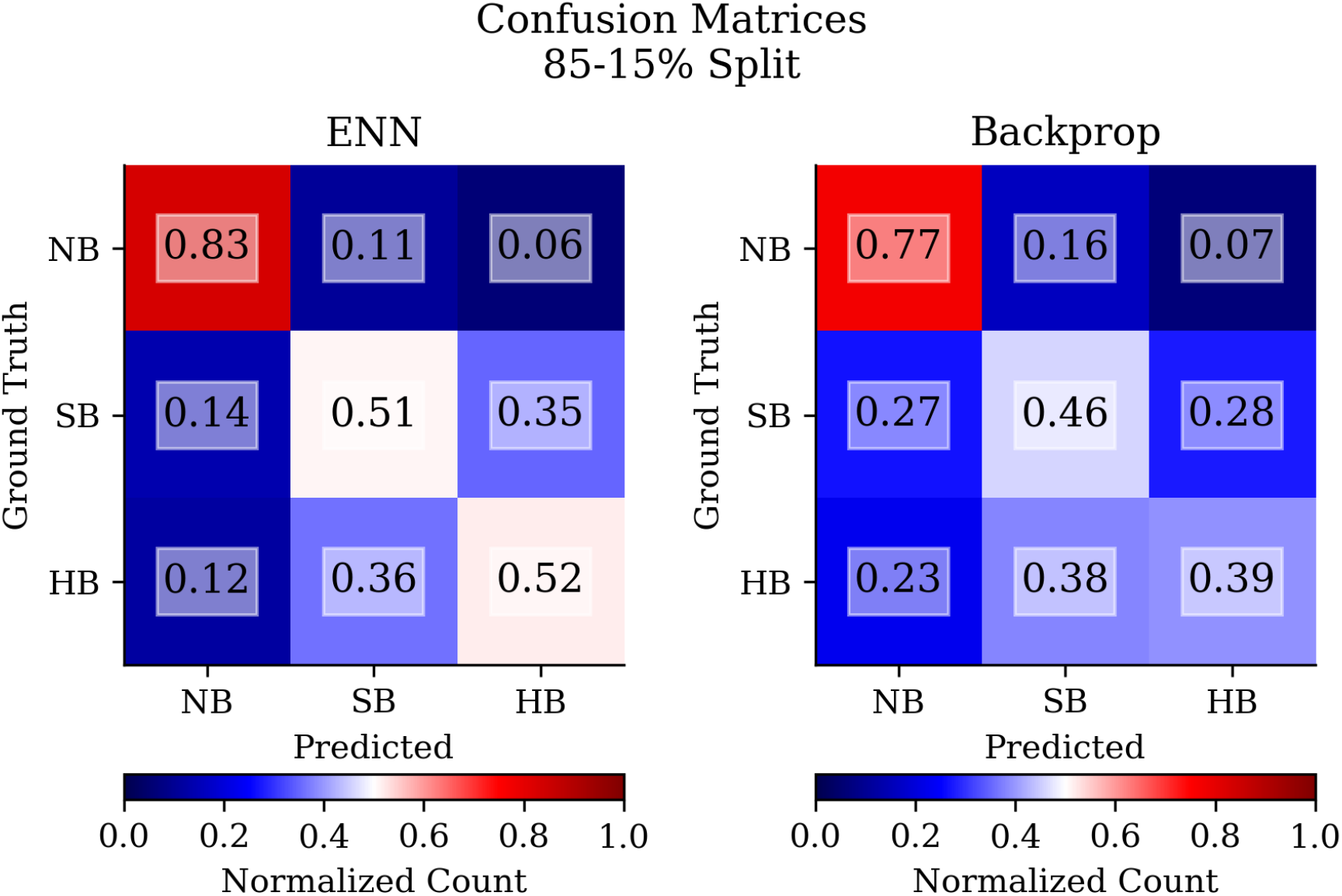
Confusion matrix of boundary-detection. Confusion matrices of network class accuracies for Backpropagation and ENNs on variable 85-15% splits of the dataset.

**Figure S3:**
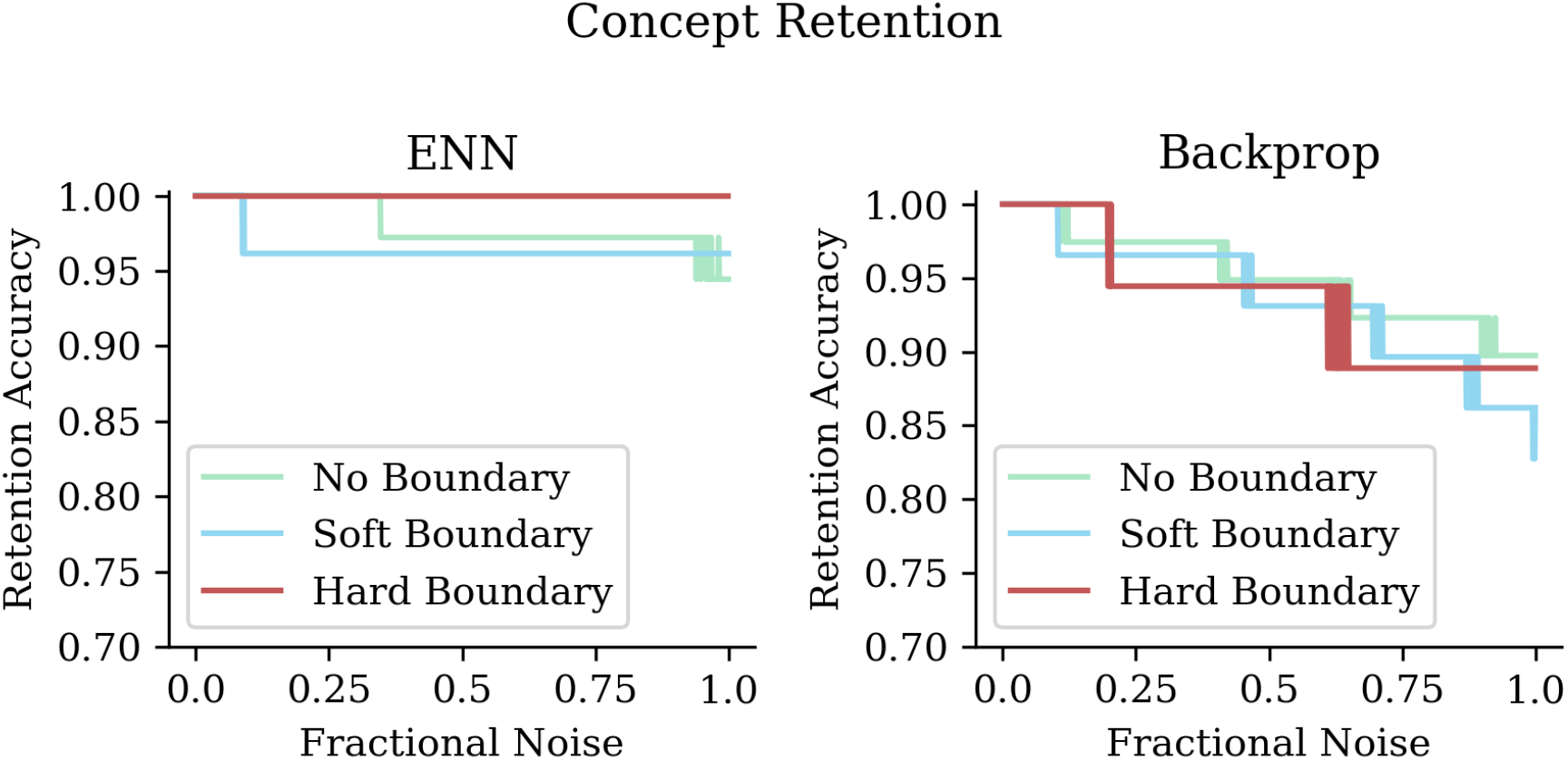
Boundary-detection retention by class under noise-injection. Network retention accuracy by class for noised networks.

**Figure S4:**
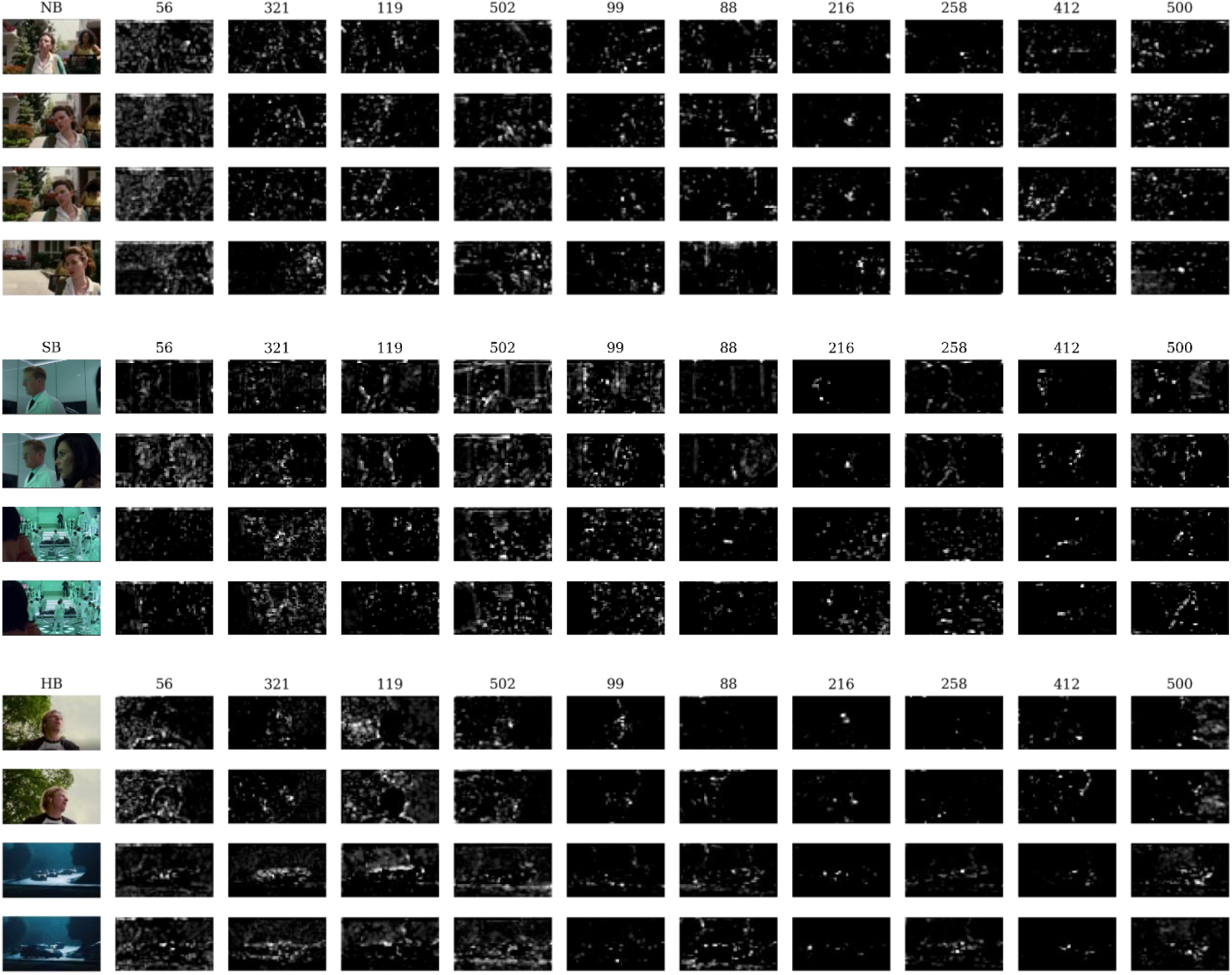
Feature maps of top ENN-derived features. Example top features from ENN-consensus and their VGG feature maps across example clips from each class.

**Figure S5:**
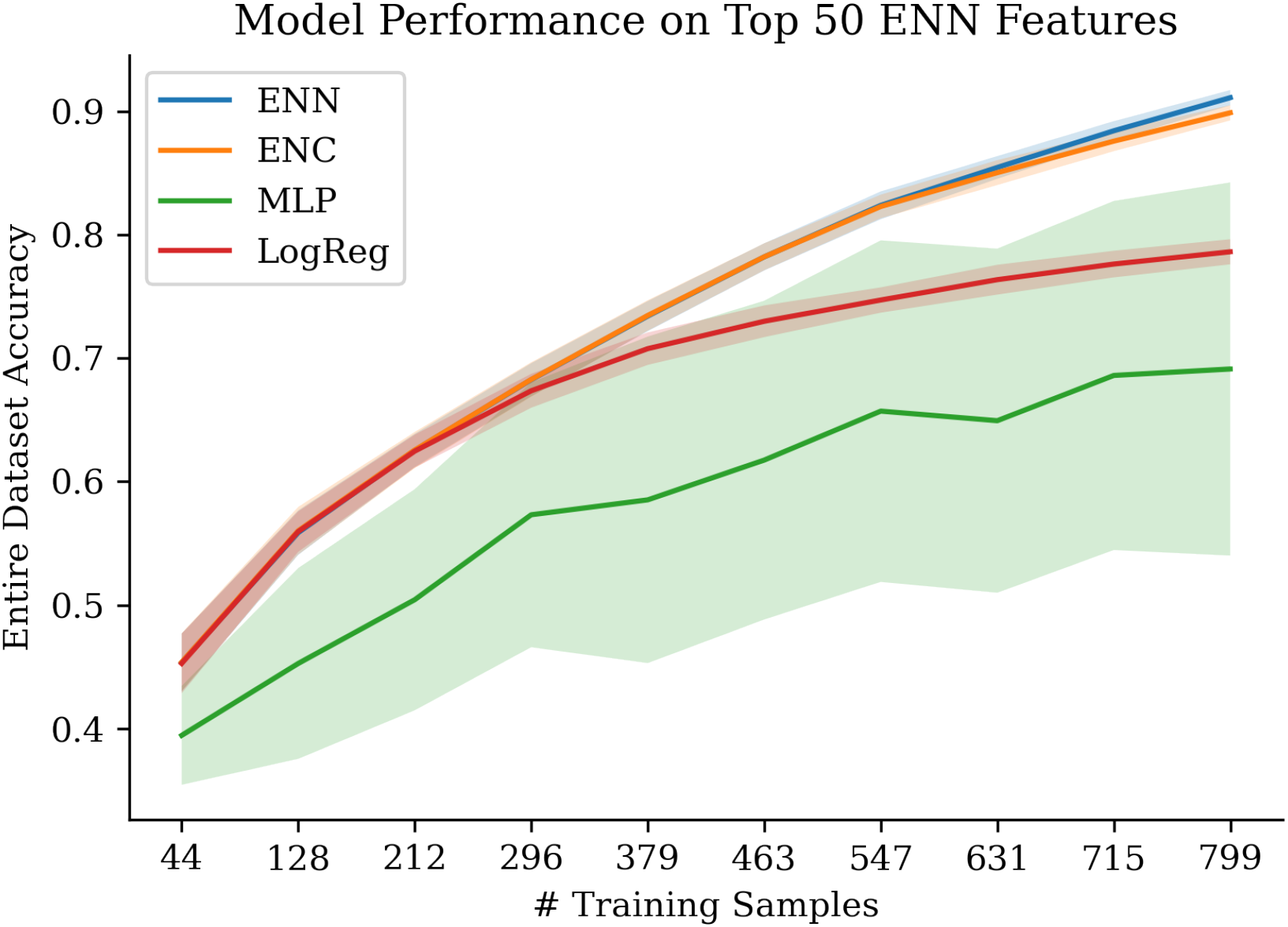
Network performance with sparsified dataset. Performance of networks and distilled circuit (ENC) when trained on a subset of the sparsified dataset (50 features derived from ENN consensus) and evaluated on the entire dataset (train and test splits). Average and standard deviation of 100 trials at 10 varying splits shown.

## Notes

### Competing Interest Statement

The authors have declared no competing interest.

